# Retinal tau phosphorylation in Alzheimer’s disease: a mass spectrometry study

**DOI:** 10.1101/2025.02.17.638688

**Authors:** Jessica Santiago, Dovile Poceviciute, Jacob Vogel, The Netherlands Brain Bank, Gunnar Brinkmalm, Malin Wennström

## Abstract

Most neurodegenerative diseases, including Alzheimer’s disease (AD) and multiple sclerosis (MS), are associated with abnormal phosphorylation of tau (p-tau) in the brain. Immunostaining studies have revealed an accumulation of p-tau in the AD retina and suggested it may reflect tau pathology in the brain. To validate these findings and further investigate the relationship between retinal and brain tau pathology, we used mass spectrometry to measure p-tau peptides in matched retinal and hippocampal samples from non-demented controls (NC, n=8), AD (n=12), and MS (n=4). We analysed the differences in p-tau levels between diagnoses and explored how retinal p-tau variants correlate with hippocampal p-tau and neuropathological changes. Our results revealed peaks corresponding to tau peptides phosphorylated at T181, S199/S202, T231, S396 + T403/S404, and T403/S404. These p-tau peptides were also detected in the hippocampal samples, along with additional p-tau peptides such as T217 and S262. Total tau phosphorylation and phosphorylation at S199/S202 and T231 were significantly higher in the retina of AD cases compared to NC. These two peptides, along with peptides phosphorylated at S396+T403/S404 and T403/S404, were also higher in cases with high amyloid-beta (Aβ) Braak stages compared to those with low Aβ Braak stages. Higher Aβ Braak stages were also associated with higher mass spectrometry peak intensities of peptides phosphorylated at S199/S202 and S396+T403/S404. Additionally, retinal p-tau peptides at S396+T403/S404 and T403/S404 correlated with neurofibrillary tangle (NFT) Braak stages, and p-tau peptides S396+T403/S404 in the retina were linked to corresponding phosphorylation in the CA1 region. These findings underscore the connection between retinal and brain tau pathology and highlight the potential of retinal tau as a biomarker for AD diagnosis and monitoring while also deepening our understanding of tauopathies in both the retina and brain.

## INTRODUCTION

Tau is a microtubule-associated protein first isolated in 1975 [58]. The human tau protein is encoded by the MAPT gene, located on chromosome 17, and is abundant in the central and peripheral nervous systems [8, 19]. Alternative splicing of the gene’s mRNA transcript results in six distinct protein isoforms present in the central nervous system, which can have either three (3R) or four (4R) microtubule-binding repeats in the carboxy-terminal region, along with zero to two (0-2N) inserts in the amino-terminal region [57]. Tau is primarily found in neurons but is also present at low levels in glial cells [8, 36]. The protein has many physiological functions. In healthy neurons, it is primarily distributed in axons, where it interacts with tubulin to promote microtubule assembly and stability, facilitating intracellular transport and neuronal signaling. To a lesser extent, tau is also localized in dendrites, where it targets factors that modulate postsynaptic receptor activity [8, 27, 28]. Additionally, some tau is found in the nucleus, where it may play a role in maintaining DNA integrity [50, 54].

Tau can undergo numerous post-translational modifications (PTMs), including phosphorylation, acetylation, ubiquitination, and methylation [2, 23, 57]. Phosphorylation of tau (p-tau) is the most common PTM and refers to the reversible addition of a phosphate at different sites of the protein [2]. The longest tau isoform in the brain (2N4R) contains up to 85 potential phosphorylation sites, many of which are accessible due to their unfolded structure [42]. In healthy brains, about 20 of these sites can be phosphorylated, which is normal and necessary for tau’s function [2, 37]. However, in individuals with neurodegenerative diseases involving tau, around 45 sites are found to be abnormally phosphorylated. Some of these sites overlap with those that are phosphorylated in healthy brains, indicating an excessive and abnormal increase in phosphorylation in these conditions [2, 23].

Abnormal PTMs can cause tau to detach from microtubules, leading to microtubule disassembly in axons. This detached tau can mislocalize into presynaptic terminals, inducing synaptic dysfunction and reducing the number of synaptic vesicles and synapses [15]. Additionally, tau may enter dendrites and postsynaptic compartments, causing postsynaptic dysfunction and synapse loss [2, 27, 51]. Pathological tau’s inability to enter the nucleus may lead to DNA damage due to the loss of its DNA-protective function [50]. Finally, tau may form aggregates that can mediate neuroinflammation and potentially induce pathological effects, leading to the deterioration of neuronal function and the development of neurodegenerative disorders. Abnormal phosphorylation disrupts tau’s normal function, leading to the formation of paired helical filaments (PHFs) and neurofibrillary tangles (NFTs), which are pathological hallmarks of tauopathies, such as Alzheimer’s disease (AD)[22]. These aggregates can be released into the extracellular space, taken up by other neurons, and spread tau pathology[12, 56].

In AD, tau pathology is crucial for diagnosis and is closely linked to cognitive decline and neuronal injury. In the brain, the disease is marked by elevated levels of hyperphosphorylated tau (p-tau), forming NFTs that are together with amyloid beta (Aβ) plaques, hallmarks of the disease [5, 24]. The severity of the tau pathology and its progression are quantified using the Braak staging system, which measures the extent of disease-related tau aggregation[9]. Abnormal tau phosphorylation is also a common feature of other neurodegenerative disorders, including multiple sclerosis (MS), where studies have identified hyperphosphorylated tau aggregates in the brain [3, 4] and suggested a role for tau pathology in driving disease progression[3, 33].

Interestingly, recent studies have detected disease-characteristic changes in the AD retina including accumulation of Aβ [1, 11, 16, 18, 21, 29–32, 35, 41, 45, 47, 48, 52, 59], islet amyloid polypeptide [46], and phosphorylated TAR DNA binding protein-43[25]. In particular, p-tau in the retina of AD patients has gained recent attention [16, 18, 25, 26, 38, 44, 49], and studies have demonstrated its correlation with tau pathology in the brain [25, 26, 38, 49]. These findings are promising as they suggest that retinal p-tau could serve as a potential non-invasive biomarker for diagnosing and monitoring the progression of AD. Histopathological analysis of postmortem retinal tissue revealed total tau (HT7, 43D), 3-repeat and 4-repeat tau isoforms (RD3, RD4), and various p-tau epitopes in retinal cross-sections from AD patients, such as p-tau at T181, S202+T205, T212+S214, T217, T231 and S396 [26, 38]. Immunoreactivity of p-tau S202+T205 and p-tau T217 were significantly elevated in the retina of AD cases compared to controls and correlated with the Braak stage for NFTs and with p-tau S202+T205 levels in the hippocampus and cortex of these patients [26, 49]. The co-localization of p-tau at sites of neuronal loss in the human AD retina [13, 21] may indicate this tissue might share a similar tauopathy with the brain.

Despite these findings, our understanding of tauopathy distribution, formation, spread, and impact on retinal inflammation and degeneration remains limited. The connections between abnormal retinal tau and brain AD pathology, as well as cognitive function, are also not completely understood. While previous studies have relied on immunohistochemistry and other antibody-based technologies to detect retinal p-tau, these methods have limitations. Variability in results can arise from differences in sample preparation, antibody specificity, and staining protocols. Advances in mass spectrometry now allow for more precise identification and quantification of tau and its modifications in retinal tissues. In this study, we aimed to investigate retinal p-tau using mass spectrometry, focusing on the specific phosphorylated peptides detected in the retina, their similarities to those found in the hippocampus, and their potential connections to AD pathology. This method enhances accuracy and yields new insights into the role of retinal tau in AD pathology.

## MATERIAL AND METHODS

### Cases included in the study

Matched retina and hippocampi were postmortem-collected from (n=8) non-demented controls (NC), (n=12) AD, and (n=4) MS cases (The Netherlands Brain Bank (NBB). For the statistical analysis of the retina, individuals diagnosed with macular degeneration or those with insufficient protein levels that could affect normalization (less than 20% of the median) were excluded. The demographics of the cases are found in Table 1 and Supplementary Table 1. The Braak staging system was used to assess the presence of both Aβ plaques and tauopathy. Aβ plaques were classified into four levels, O, A, B, and C, representing none, some, moderate, and many plaques, respectively. The presence of tauopathy in the brain was scored into stages I–VI based on their distribution. NFTs were observed in the entorhinal cortex and hippocampus in stages I and II, while stages III and IV indicated NFT spread to the temporal neocortex, and stages V and VI showed widespread distribution throughout the brain [10]. Written consent was obtained from all patients or their relatives for the use of tissue and clinical data in research, in accordance with the international declaration of Helsinki and Europe’s code of conduct for Brain Banking. The collection procedures were approved by the medical ethical committee of the VU Medical Center Amsterdam, and the studies were approved by the regional ethical review board in Lund.

**Table 1.** Cases included in the study.

|  | NC | AD | MS |
| --- | --- | --- | --- |
| Number | 8 | 12 | 4 |
| Age (mean ± SD) | 79.0 ± 12.8 | 80.7 ± 10.9 | 76.5 ± 10.1 |
| Females (%) | 50.0% | 75.0% | 75.0% |
| APOE4 positive (%) | 37.5% | 66.7% | 25.0% |
| PMD (mean ± SD) | 5:49 ± 1:10 | 6:10 ± 1:21 | 8:49 ± 0:52 |

NC=non demented controls, AD=Alzheimers disease, MS=Multiple sclerosis, APOE4=apolipoprotein E4 gene, PMD=postmortem delay

### Dissection of retina and hippocampi samples and preparation for mass spectrometry

At the autopsy, the lenses of the right eye of the cases were removed and filled with O.C.T. mounting media (Vector laboratories) and thereafter frozen and stored at −80 °C. The eyeball was cut into eight clefts, sparing 0.5 cm around the optic nerve. The retina was thereafter collected from the middle periphery region of the cleft from the nasal superior region. The hippocampi were snap-frozen at autopsy in one-centimetre-thick sections and stored at −80 °C. For analysis, the samples were divided into two half centimetre-thick sections. A 3 mm biopsy punch (Kai Medical) was used to dissect a sample from the Cornu Ammonis 1 (CA1) region next to the Subiculum from one of the sections. The CA1 was chosen as it starts to exhibit NFTs and Aβ plaques during the early/moderate stages of AD [10]. The retina and hippocampi samples were transferred to 1.5 mL Pink Rino Tubes with screw caps and 100 µL Lysis buffer (50 mM Tris-HCl pH 7.5, 50 mM NaCl, 1 mM EDTA, 5 mM NaH2PO4, 1 mM DTT) was added and placed in the Bullet Blender Storm Pro (BT24M, Next Advance, Inc. Troy, NY, USA). The samples were run for 3 min at speed 8 and thereafter centrifuged at 14 000 x g for 10 min. The supernatants were transferred to new Eppendorf tubes. The bullets were washed with 50 µL Lysis buffer followed by centrifugation, and the supernatants were added to the Eppendorf tubes. The remaining material in the Rino Tubes was subjected to a new round of sample preparation now using Ripa-buffer (50 mM Tris-HCl pH 7.4, 150 mM NaCl, 1mM EDTA, 1% Triton X-100, 0.1% sodium deoxycholate) instead and collected into new Eppendorf tubes.

From each sample, 50 µL was taken and reduced with DTT to a final concentration of 10 mM at 56 °C for 30 min, followed by alkylation with iodoacetic acid to a final concentration of 20 mM for 30 min in the dark. The samples were precipitated with ice-cold ethanol (final concentration of 90% ethanol) overnight at −20 °C. Samples were centrifuged at 14 000 x g for 10 min. The pellets were allowed to airdry and resolved in 50 µL 100 mM ammonium bicarbonate, and the samples were disrupted using a BioRuptor (Diagenode Inc., Denville, USA), with the following settings: 20 cycles, each consisting of 15 sec on/off. Samples were centrifuged at 14 000 x g for 10 min, supernatants were transferred to new tubes, and the protein concentration was determined by NanoDrop (DeNovix, AH diagnostics) at A280 nm. The samples, 15 µg, were digested with trypsin (Promega, Madison, WI) in a ratio of 1:50 w/w (enzyme: proteins) overnight at 37 °C. The digestion was stopped by 5 µL 10% trifluoroacetic acid (TFA). The samples were dried using a Speed Vac and resolved 22 µL in 2% ACN/0.1% TFA.

### LC-MS/MS analysis

The samples, 2 µL, were analysed on an Exploris 480 mass spectrometer (Thermo Fischer Scientific) coupled with a Vanquish Neo UHPLC system (Thermo Fischer Scientific). Two-column setup was used on the HPLC system, and peptides were loaded into an Acclaim PepMap 100 C18 precolumn (75 μm x 2 cm, Thermo Scientific, Waltham, MA) and then separated on an EASY spray column (75 μm x 25 cm, C18, 2 μm, 100 Å, ES902) with the flow rate of 300 nL/min. The column temperature was set at 45 °C. Solvent A (0.1% FA in water) and solvent B (0.1% FA in 80% ACN) were used to create a 120 min nonlinear gradient from 5 to 25% of solvent B for 100 min and increased 32 [25]% for 12 min and the increased to 45% for 8 min to elute the peptides.

The samples were analysed with a data-dependent acquisition (DDA) in positive mode. The full MS1 resolution was set to 120 000 at m/z 200, and the normalized AGC target was set to 300% with a maximum injection time of 45 ms. The full mass range was set at 350-1400 m/z. Precursors were isolated with the isolation window of 1.3 m/z and fragmented by HCD with the normalized collision energy of 30. MS2 was detected in the Orbitrap with a resolution of 15,000. The normalized AGC target and the maximum injection time were set to 100% and custom, respectively. The intensity threshold for precursor selection was set to 104, and 60 s dynamic exclusion was applied.

The generated mass spectra were analysed using Proteome Discoverer 2.5 (Thermo Fisher Scientific) and PEAKS (version Xpro) against the UniProt Human database (UP000005640, canonical) to identify the tau phospho-peptides and the most probable location of the phosphorylations. The precursor tolerance and fragment tolerance were set to 10 ppm and 0.02 Da, respectively. Trypsin was selected as the enzyme, methionine oxidation, deamidation of asparagine, phosphorylation, oxidation, and acetylation were treated as dynamic modifications, and carbamidomethylation of cysteine was treated as a fixed modification. The peak intensities from Proteome Discoverer were used for statistical analysis by label-free relative quantification. Peak heights were used for intensity determination, and the extracted chromatographic intensities were used to compare peptide peak intensities across samples.

PEAKS Studio (version Xpro) was used to evaluate the details of tau PTMs. Searches were made against a custom-made database containing only tau isoforms with phosphorylation (STY), oxidation (M), and acetylation (protein N-term) as variable modifications. Search tolerances were 20 ppm and 0.05 Da for precursor and fragment masses, respectively, and enzyme specificity was set to unspecific. In addition, a PTM search was performed with 313 preselected PTMs, including methylation, acetylation, and ubiquitination at relevant amino acids. MS/MS data was manually inspected to validate the positions of phosphate groups. When the precise location was not possible to determine, or both variants were present, this is indicated (e.g., p199/202). An overview of the workflow for sample selection, preparation, and analysis is shown in Figure 1. Retinal and hippocampal tissues underwent protein extraction, were digested into peptides, and analysed using Liquid Chromatography-Tandem Mass Spectrometry (LC-MS/MS). Data were processed for peptide identification, quantification, and statistical analysis.

**Figure 1.**
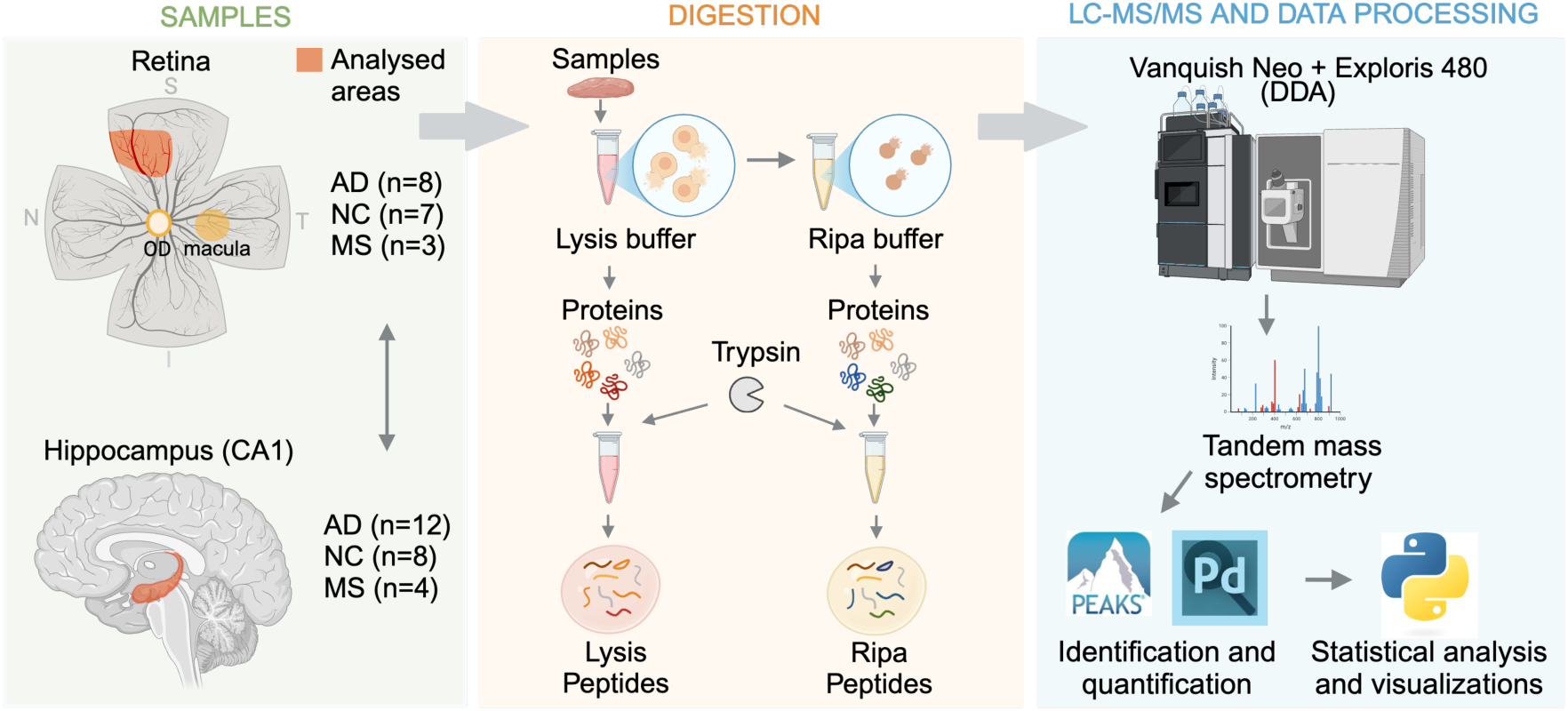
Overview of sample preparation, digestion, and data processing workflow. Retinal (middle nasal superior area) and hippocampal (CA1 area) tissues were collected from Alzheimer’s disease (AD), non-demented control (NC), and multiple sclerosis (MS) cases. The retina’s anatomical orientation is as follows: I (inferior), S (superior), N (nasal), T (temporal), and OD (optic disc). Protein extraction was carried out sequentially using a Lysis buffer to target extracellular components and the cytoplasm, followed by a Ripa buffer to extract proteins from nuclei and organelles. The resulting extracts were digested with trypsin to generate proteolytic peptides. Peptides were analysed using the Vanquish Neo + Exploris 480 Liquid Chromatography-Tandem Mass Spectrometry (LC-MS/MS) system in data-dependent acquisition (DDA) mode. Data were processed for peptide identification and quantification using Proteome Discoverer (PD) and PEAKS software, with statistical analysis and visualizations conducted in Python. Created in BioRender. Santiago, J. (2025) https://BioRender.com/e79t121.

### Immunofluorescence staining procedures

Flat mounts of the far periphery of the nasal superior retina of the right eye from (n=1) AD case and (n=1) NC were left in 4% paraformaldehyde for 4 h and thereafter rinsed in PBS and stored in antifreeze −20 °C until staining. The second half-centimeter-thick hippocampal sample of one AD case was left in 4% paraformaldehyde for 4 h and, thereafter, in 30% sucrose for 2 days. Thereafter, the sample was sectioned into 40 µm thick free-floating sections and stored in antifreeze −20 °C until staining.

The retina flat mounts and hippocampi sections were stained according to the following procedures: Samples were washed with phosphate-buffered saline (PBS) three times and incubated in blocking solution (BS) consisting of 5% goat serum and 0.25% triton in PBS for 1 h at room temperature (RT). Thereafter, primary antibodies (Table 2) were added to the BS, and the samples were left over three nights at 4°C on a slow shaker. After the third night, the samples were rinsed 6 times in PBS+0.25% triton and subsequently incubated with the appropriate secondary antibody (Goat anti-Rabbit 488 or Goat anti-Mouse 488 from Invitrogen) for 2 h in RT. Finally, the samples were mounted on glass slides covered with. The hippocampi sections were incubated in Sudan Black (1% in 70% ethanol) (Sigma-Aldrich) for 5 min before they together with the retinal flat mounts, were mounted with Vectashield Set mounting medium containing DAPI (Vector Laboratories). Negative control for each retinal staining was included in the immunostaining procedure by omitting the primary antibody, thereby confirming the specificity of the observed signal and ruling out nonspecific binding of the secondary antibody or background staining. The immunostainings of hippocampi samples served as positive controls. Images of the immunostainings of retina were captured using a confocal microscopy (Zeiss LCM 800) with the 63x objective, while the images of the brain immunostainings were captured using an Olympus AX70 light microscope with the 20x objective.

**Table 2.** Primary Antibodies used in the study.

| Antibody | Target | Isotype | Dilution | Cat.no | Source |
| --- | --- | --- | --- | --- | --- |
| p-Tau | T181 | Rabbit IgG | 1:200 | Sc-101816 | Santa Cruz |
| CP13 | S202 | Mouse IgG | 1:200 | N/A | Dr Peter Davies |
| ERP2488 | T231 | Rabbit IgG | 1:200 | Ab151559 | Abcam |
| PHF-1 | S396+S404 | Mouse IgG | 1:200 | N/A | Dr Peter Davies |
| AB_253376 | S404 | Rabbit IgG | 1:200 | 44-758G | Invitrogen |

### Statistical Analysis

All analyses were carried out using Python (v.3.9.12) with the scipy.stats [55] and statsmodels. Data visualizations were generated using the matplotlib and seaborn libraries. The sample size was not predetermined by statistical methods; instead, all available data were included in the analysis. All tau phospho-peptides identified in the tissue were described for comparisons between groups and correlations. p-tau peptides and subjects with more than 50% of missing values were not included in the statical analysis. For the remaining, missing values were imputed with a value corresponding to 50% of the minimum observed value for the respective peptide. Mass spectrometry peak intensity values were normalized to the median and log-transformed prior to analysis. Subjects with any preparation showing less than 20% of the median protein values were excluded, as the required correction factor would compromise data reliability. Comparisons were conducted using mixed effects models with peptide abundance as the dependent variable, diagnosis or Aβ high/low classification and buffer fraction, as well as their interaction, as fixed effects, and fitting random intercepts for donor. Following this, we used visualization and t-tests as posthoc tests to interpret the direction of the interactions. Correlations were evaluated using Spearman’s correlation tests. All statistical tests were two-sided, with a significance threshold of p < 0.05. Multiple testing correction using Benjamin-Hochberg was applied when applicable, and the adjusted results were reported with a significance threshold of a FDR-adjusted p-value (q-value) < 0.05.

## RESULTS

### Phosphorylation sites of tau protein in retina and CA1

The first step of the study was to identify tau phosphorylation (p-tau) sites in the retina and the brain of AD, NC, and MS cases. For this analysis, the samples were homogenized and sequentially treated with two distinct buffers (see also Figure 1). First, a detergent-free Lysis buffer was used to isolate soluble cytoplasmic and extracellular components. Thereafter, the pellet was homogenized and dissolved in a detergent containing Ripa buffer to extract insoluble proteins and nuclear and organelle-associated structures. The findings, summarized in Table 3, show the peptides with p-tau sites found in the retina and CA1 of the hippocampus under the two different buffer conditions. In the retina, tau peptides with phosphorylation at T181, S199/S202, T231, T231+T235, S396 + T403/S404, and T403/S404 were observed in both Lysis and Ripa buffers. The p-tau peptides found in the CA1 region included those observed in the retina, as well as additional unique p-tau peptides. For instance, tau peptides phosphorylated at the sites S205, T212/S214, T212+T217, T217, S262, S396 + S400 + T403/S404 were consistently observed across both CA1 buffer conditions. Peptides with phosphorylation at the sites S356 and S396 were detected in the CA1 Lysis buffer but not in the CA1 Ripa buffer. The found phosphorylation sites in the retina were also evaluated by staining one NC and one AD flat mount sample against T181, S202, T231, S396 + S404, and S404 (Figure 2). Staining against all p-tau sites in NC gave rise to a dotted pattern (Figure 2A), while the staining of the AD sample instead revealed a stripped pattern (Figure 2A), resembling neuronal processes. The positive control, i.e., the hippocampal section from an AD patient, showed the expected tangles and neuropil threads positive for T181, S202, T231, T231, S396 + S404, and S404 (Figure 2C).

**Figure 2.**
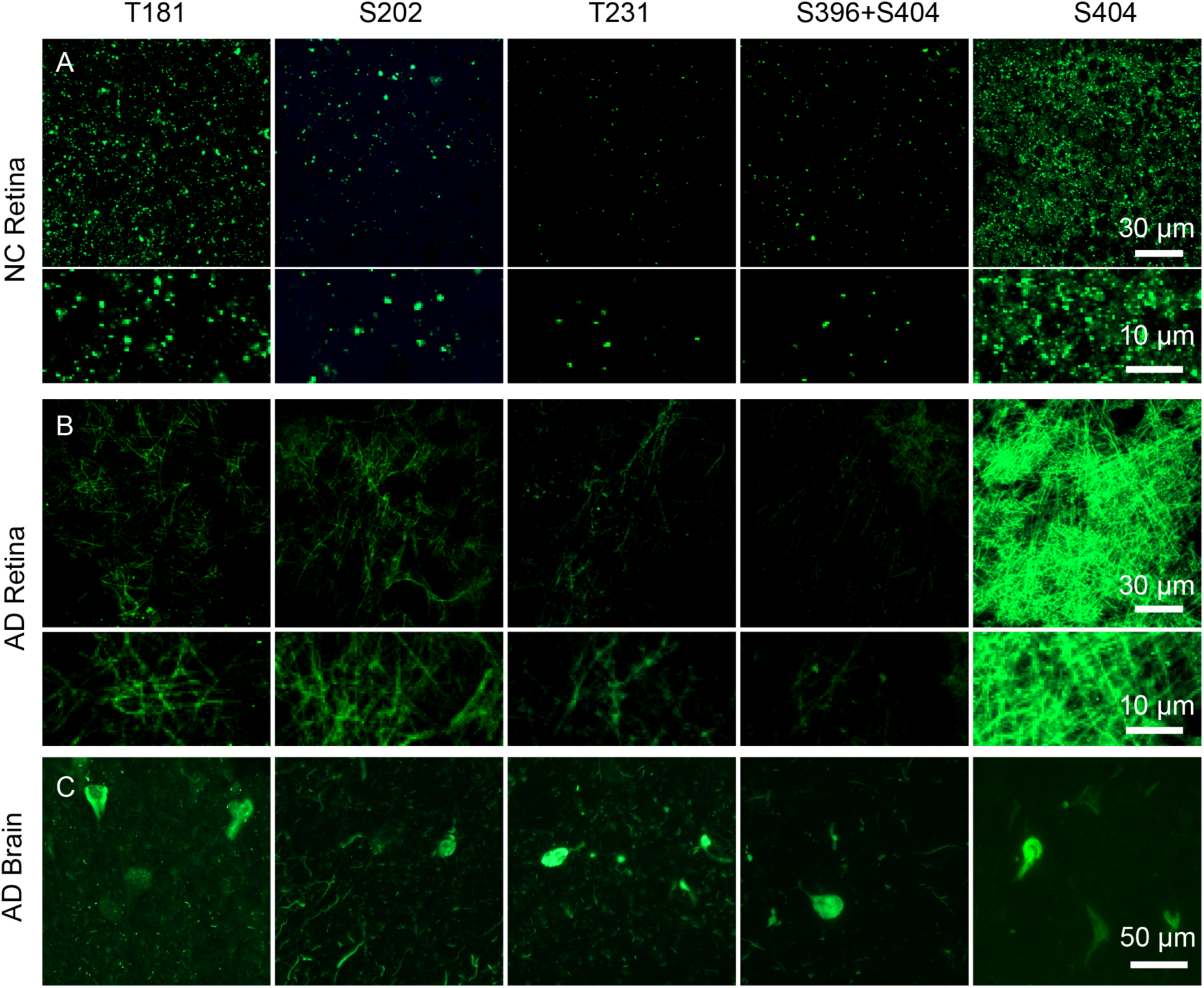
Immunostainings against phosphorylated tau epitopes in flatmounts of retina. Images in (A) show the dotted pattern found in immunostainings of retinal flatmounts from a non-demented control (NC) stained against T181, S202, T231, S396+S404, and S404. The images were captured with a confocal microscope (x63 objective), and the upper image shows an overview of the staining, while the lower image shows an enhanced view of the staining. Images in (B) show the overview and enhanced view of the stripped pattern revealed in the immunostainings of retinal flatmounts from a case with Alzheimer’s disease (AD) stained against T181, S202, T231, S396+S404, and S404. Images in (C) show the characteristic tangles and neuropil threads in the positive control (AD brain), detected by the T181, S202, T231, S396+S404, and S404 antibodies used in the study.

**Table 3:**
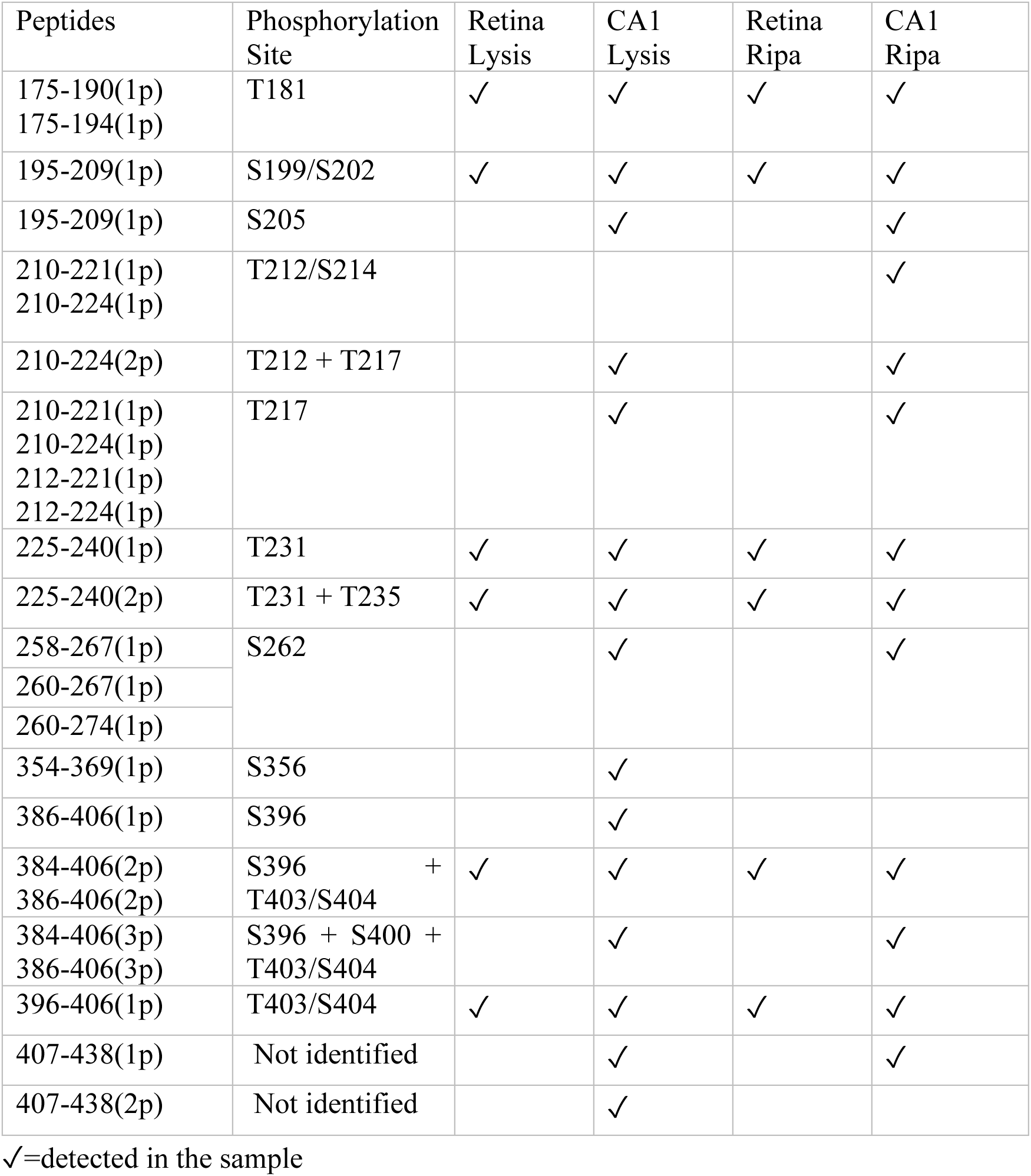
Summary of p-tau peptides identified in the retina and CA1 of the hippocampus after homogenization in Lysis and Ripa buffers.

### Relative distribution of p-tau peptide mass spectrometry peak intensity in retina and CA1

Next, we analysed the relative distribution of the peak intensities of phosphorylated tau (p-tau) peptides in the retina and CA1 across different buffer conditions and diagnostic groups (NC, MS, and AD). For this analysis, only peptides detected in more than 50% of samples were considered. The results revealed that the peptide with p-tau at T403/S404 yielded the most intense peaks in the retina and CA1 regardless of diagnosis or buffer condition. Peptides with phosphorylation at T181 and S199/S202 exhibited high signals in both tissues and buffer conditions and all diagnosis groups. In the brain, the S199/S202 peptide signals were slightly higher than the T181 peptide, while the opposite was observed in the retina, although the signals were very close in this tissue. Interestingly, peptides with phosphorylation at T231 showed distinct patterns, with high signals in the retina Lysis buffer, and lower peaks in both buffers in CA1. Notably, in the retina Ripa buffer, the p-tau T231 peptide intensities were higher in NC and MS than in AD. The signal intensities of the peptide with double phosphorylation T31+T235 were low in all preparations and showed different values across diagnosis in retina Ripa buffer, with higher intensities in MS and NC than in AD. Finally, peptides with phosphorylation at S396+ T403/S404 displayed the lowest intensities in the retina and CA1 regardless of diagnosis and buffer type (Figure 3).

**Figure 3.**
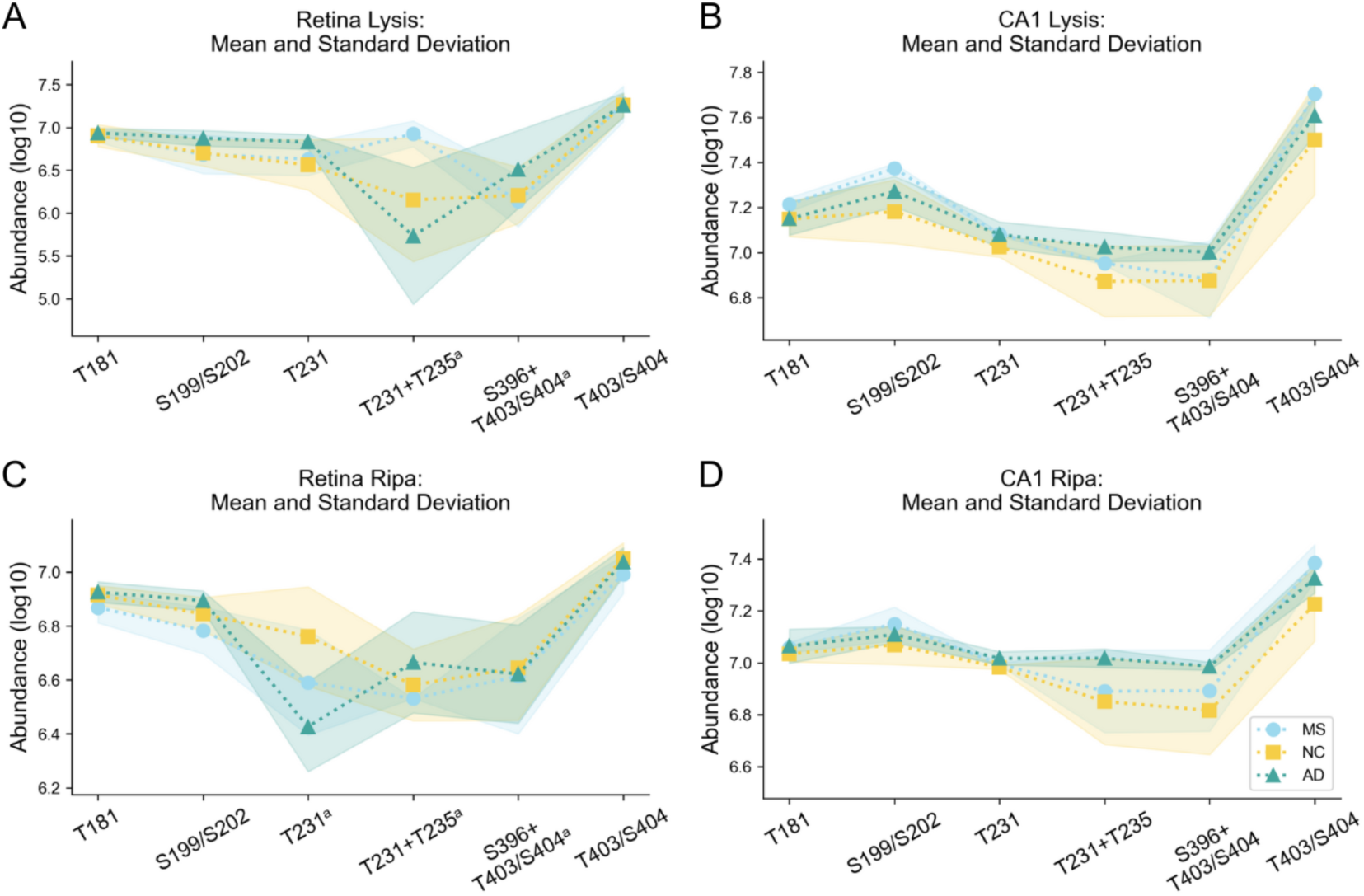
The relative distribution of tau peptides phosphorylated at different sites in retina and CA1. Peak intensities of six tau peptides phosphorylated at T181, S199/202, T231, T231+T335, S396+T403/S404 and T403/404 in retina and the cornu Ammonis 1 (CA1) of hippocampus. The graph in (A and B) shows the mean normalized signal of phosphorylated tau peptides in retinal (A) and CA1 (B) Lysis samples from NC (yellow), MS (blue), and AD (green). The graph in (C and D) illustrates the peak intensity values in the Ripa extraction of retina (C) and CA1 (D) of NC (yellow), MS (turquoise), and AD cases (green). Missing values were replaced by an imputed value representing 50% of the lowest value, p-tau peptides marked with an (ᵃ) were detected in less than 50% of the subjects. Shaded regions indicate standard deviation.

### Differences in peak intensities of p-tau peptides between diagnosis groups

To investigate if the presence of the phosphorylated peptides differed between groups we compared the peak intensity values in NC, AD and MS. A mixed effects model was applied to evaluate the impact of the three diagnoses, the two buffer fractions (Lysis and Ripa), and their interaction on the presence of phosphorylated peptides in the retina. The results revealed a significant buffer fraction by diagnosis interaction for peptides phosphorylated at S202 (t=−2.207, p=0.027), T231 (t=−6.006, p<0.001), and the total value of phosphorylated tau peptides (i.e., the summation of the peak intensities of all p-tau peptides) (t=−3.387, p = 0.001). Specifically, the differences were more pronounced in Lysis than in Ripa. For the peptides phosphorylated at S202 (t=3.112, p=0.002), T231 (t=2.680, p= 0.007), and total p-tau (t=3.563, p<0.001), we also noted a consistent main effect of diagnosis, where AD had greater peptide abundance than NC irrespective of buffer fraction. To visualize and understand the direction of the interaction, we next analysed the data using t-tests. In AD, retinal p-tau peptides showed significantly higher intensities of total phosphorylation, phosphorylation at S199/S202, and T231 in Lysis buffer compared to NC (p=0.026, p= 0.017 and p= 0.028, respectively) (Figure 4A-C). AD also showed higher peak intensity values for peptides with p-tau T231 in Lysis buffer and p-tau S199/S202 in Ripa buffer compared with MS (p=0.034, p=0.031) (Figure 4D). To enhance the power of the analysis, we stratified the cases into high and low Aβ pathology based on Aβ Braak stages, where Aβ low represented cases with Braak stages O and A and Aβ high represented cases with Braak stages B and C. Analysis after this stratification using a mixed effects model with Aβhigh/low and buffer fraction as predicting variables revealed a significant buffer fraction by diagnosis interaction for peptides phosphorylated at T231 (t=−3.449, p=0.001) and S396+T403/S404 total (t=−2.092, p=0.036) and a greater differences in Lysis compared to Lysis. For peptides phosphorylated at S202 (t=3.112, p=0.002), T231 (t=2.860, p= 0.004), and total p-tau (t=3.529, p<0.001), as well as and S396+T403/S404 total(t, p<0.001), we also observed a consistent main effect of Aβ group, with A βhigh showing higher peptide values than Aβ low, regardless of the buffer fraction. Post-hoc t-tests showed significantly higher peak intensities of total phosphorylation of tau (p=0.038), and p-tau peptides S199/S202 (p = 0.028), T231 (p = 0.006), T403/S404 total (p=0.044) and S396+T403/S404 total (p<0.001) in the Lysis buffer, as well as S199/S202 (p=0.003) in the Ripa buffer, in the Aβ high group compared to the Aβ low group (Figure 4E-J).

**Figure 4.**
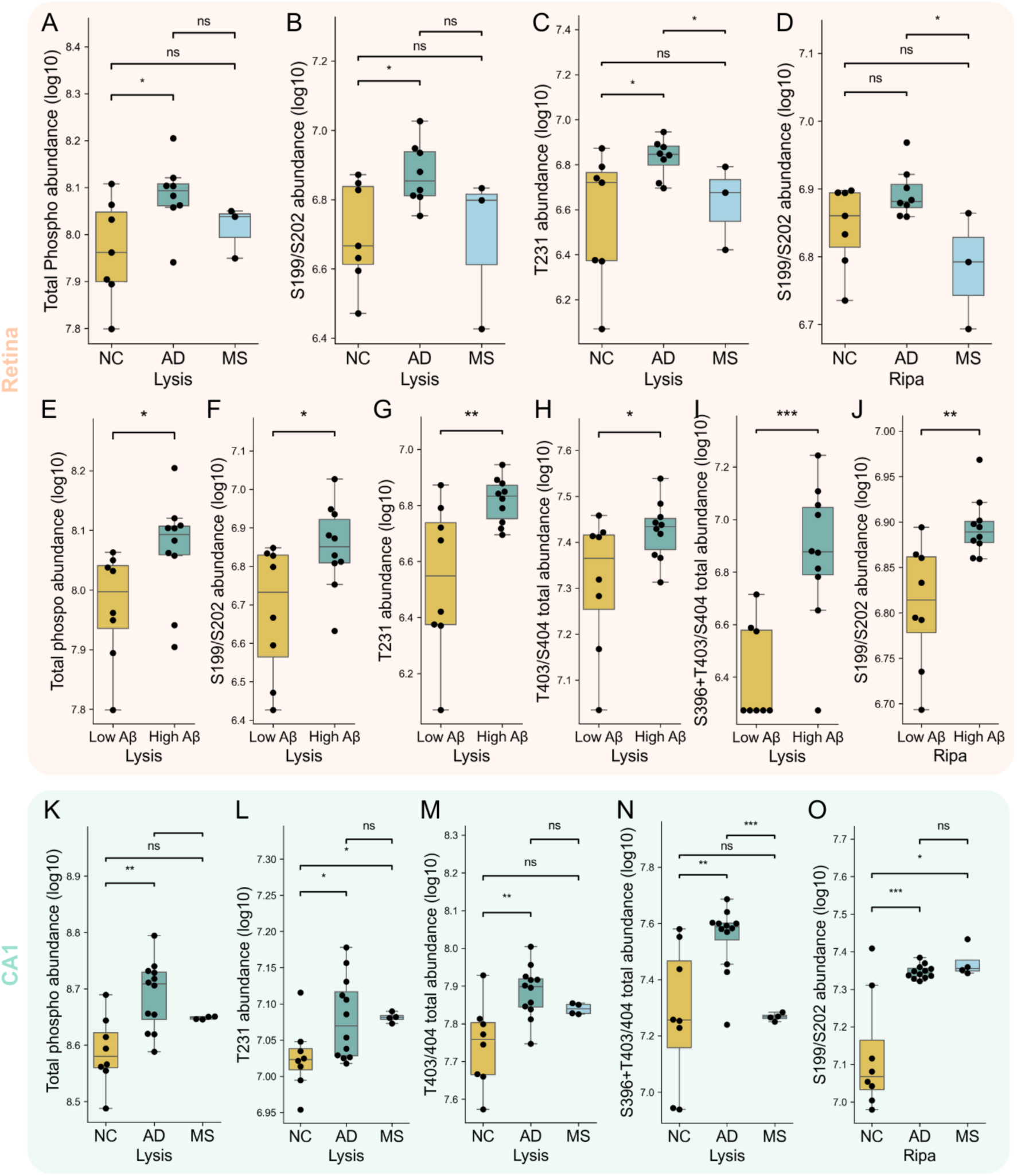
Alterations in p-tau peptides peak intensity in retinal and hippocampal tissues between stratified groups. Box plots displaying log-transformed peak intensity values of tau peptides with phosphorylation sites in retina (A–J) and CA1 (K–N). The box plot in (A) shows a higher peak intensity of peptides with total phosphorylation of tau in retina Lysis buffer from Alzheimer’s Disease (AD, green) cases compared to non-demented control cases (NC, yellow). (B) shows higher peak intensity of peptides with p-tau S199/S202 in retina Lysis buffer from AD cases compared to NC. The plot in (C) shows a higher peak intensity of peptides with p-tauT231 in retinal Lysis buffer from AD cases compared to NC and Multiple Sclerosis (MS, blue). In (D), the higher peak intensity of peptides with p-tau S199/S202 in retina Ripa buffer in MS compared to AD cases is demonstrated. Panels in (E-I) show the higher peak intensity of peptides with total p-tau, p-tau at S199/S202, T231, T403/S404, and S396+T403/S404 in Lysis buffer and at (J) S199/S202 in Ripa buffer, from cases with high Aβ compared to cases with low Aβ. The panels in (K-O) demonstrate higher peak intensity of total p-tau peptides and peptides with p-tau at S199/S202, T231, T403/S404 total and S396+ T403/S404 in CA1 Lysis buffer and at S199/S202 in Ripa buffer from AD cases compared to NC. Total refers to the sum of the peak intensity of all peptides with a given phosphorylation site, including double and triple phosphorylation. Statistical analyses were performed using t-tests. Statistical significance is indicated by asterisks (* p<0.05, ** p<0.01, *** p<0.001), and “ns” denotes non-significant differences. Error bars represent standard deviations.

To investigate if the changes in p-tau peak intensities in the retina correspond to similar changes in the brain of the same subjects, we examined whether the peptides that showed significant group differences in the retina also exhibited differences in the CA1 region. In the CA1, the results using a model considering diagnosis and buffer fractions revealed significant buffer fraction by diagnosis interaction in peak intensity between AD and NC for total T403/S404 (t=2.537, p=0.011) and total p-tau (t=2.694, p=0.007). In CA1, different from the retina, the peptides’ differences were more pronounced in Ripa than in Lysis. We also observed a main effect of AD diagnosis over NC for the peptides T231 (t=2.152, p=0.031), total S396+T403/S404 total (t=2.196, p=0.028), T403/S404 total (t=2.137, p=0.033) and total p-tau (t=2.567, p=0.010). The post-hoc t-tests showed that AD cases presented higher peak intensity of total p-tau (p=0.002), and tau peptides with phosphorylation at T231 (p=0.006), T403/S404 total (p=0.013), and S396+T403/S404 total (p= 0.004) in the Lysis buffer (Figure 4K-N). Significantly higher levels of S396+T403/S404 total were also observed in AD if compared to MS (p<0.001) (Figure 4N). The AD group also demonstrated significantly higher peak intensity of p-tau S199/S202 peptides in the Ripa buffer (p=0.001) compared to NC (Figure 4O). Additional differences in p-tau peptides between AD and NC and high vs low Aβ groups in CA1 are listed in Supplementary Tables 2 and 3. No differences in any of the phosphorylated tau peptides were observed between the sex groups.

### Correlations between retinal p-tau peptides, AD pathology, and p-tau variants in CA1

Next, we investigated if the mass spectrometry peak intensities of the different p-tau peptides in the retina correlated with the Braak stages of NFT, Lewy bodies (LB), and Aβ. The peak intensities of peptides with p-tau at T403/S404 in retina homogenized with Lysis buffer correlated positively with NFT Braak stages (r=0.47, p=0.046) (Figure 5A), and the peak intensities of peptides with p-tau at T396+T403/S404 total correlated positively with NFT, LB, and Aβ Braak stages (r=0.54, p=0.02; r=0.49, p=0.044, and r=0.76, p<0.001) (Figure 5B–D). The peak intensity of p-tau S199/S202 also correlated positively with NFT stages (r=0.51, p=0.03) (Figure 5E). We further analysed the correlation between the retinal p-tau and CA1 p-tau values but found that only S396+T403/S404 total in Lysis buffer correlated significantly between the retina and the CA1 (r= 0.515, p=0.049) (Figure 5F). Interestingly, the peak values of S396+T403/S404 total in Lysis also correlated with many other phosphorylated peptides in CA1, including p-tau at S262, T231, T217 and total phosphorylation in the Ripa buffer (Figure 5G). No correlations were found between age and any of the phosphorylated tau peptide peak intensities.

**Figure 5.**
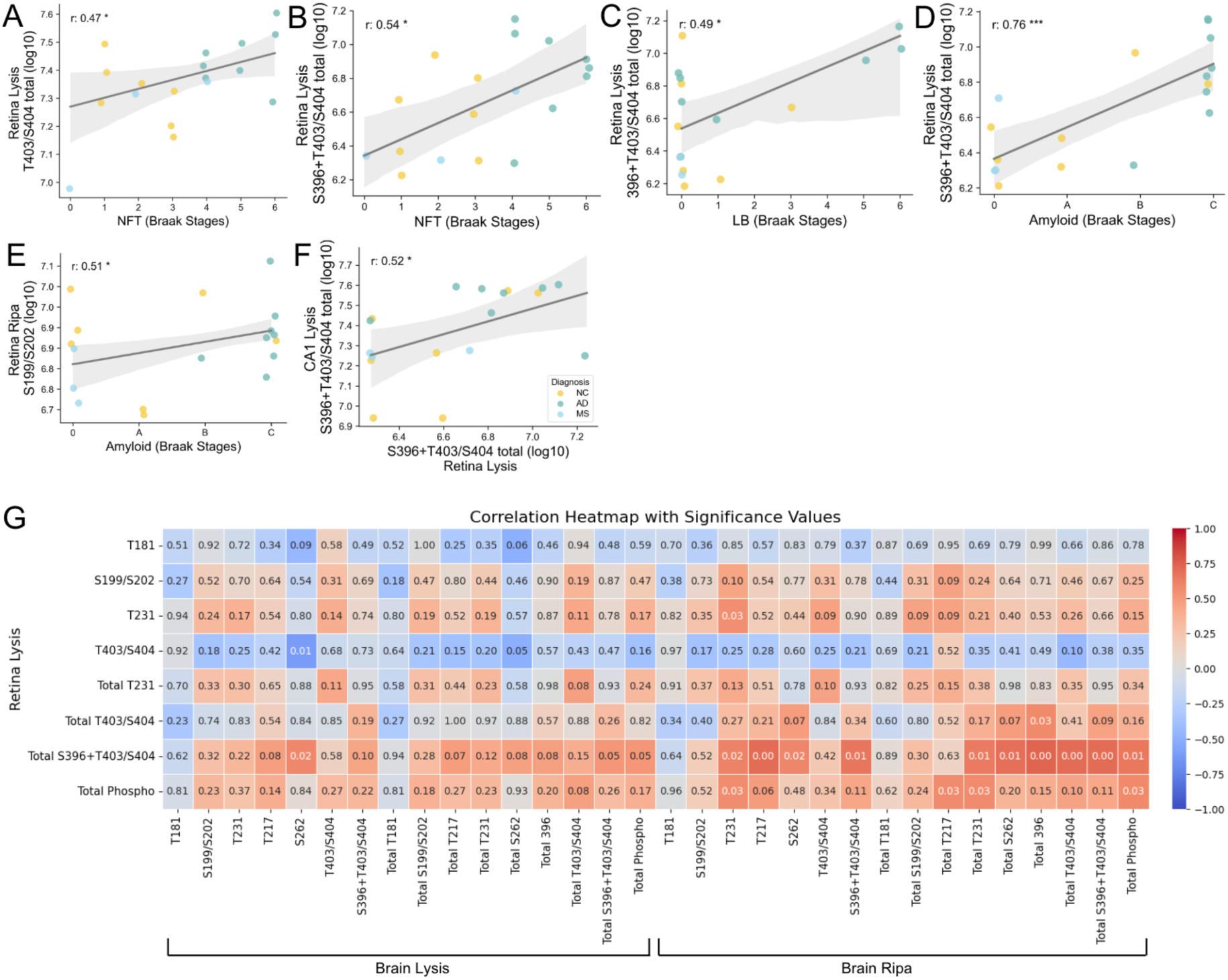
Correlation analysis of retinal p-tau peptides, AD pathology, and CA1 p-tau peptides. Scatter plots showing positive correlations between peptides with p-tau at T403/S404 total levels in retina Lysis buffer and neurofibrillary tangles (NFT) Braak stages (A), retina Lysis buffer p-tau at S396+ pT403/S404 with NFT, Lewy body (LB), and amyloid beta (Aβ) Braak stages (B, C, and D). In the Ripa buffer p-tau at S199/S202 correlated with Aβ Braak stages (E). A significant positive correlation was also found between S396+T403/S404 in the retina and CA1 Lysis buffer (F). Addtionaly, S396+T403/S404 correlated with other p-sites in the brain (G). Statistical analyses were performed using Spearman’s correlation tests. Each point represents an individual sample. Statistical significance is indicated by asterisks (* p<0.05, ** p<0.01, *** p<0.001) or shown in white in the heatmap. The colors in the heatmap correspond to r correlation values, as indicated in the color scale on the right.

### Correlations between phosphorylated tau peptides in the Retina

To assess the relationships between retinal p-tau peptides’ peak intensities, we performed correlation analyses across the peptides with the different phosphorylation sites measured in both Lysis and Ripa buffer fractions. The peak intensities of tau peptides with phosphorylation at S199/S202 and T403/S404 in Lysis buffer correlated positively with the intensities of the respective p-tau peptides in Ripa buffer (r = 0.601, p = 0.008 and r = 0.728, p < 0.001, respectively) (Figure 6A and B). Analysis of the p-tau peptides within the Ripa buffer showed significant correlations between p-tau peptides phosphorylated at T181 and S199/S202 (r = 0.669, p= 0.002; Figure 6C), as well as between p-tau peptides phosphorylated at T181 and T403/S404 (r = 0.678, p = 0.002; Figure 6D). Within the Lysis buffer, peptides phosphorylated at S199/S202 strongly correlated with p-tau T231 (r = 0.823, p < 0.001) (Figure 5E).

**Figure 6.**
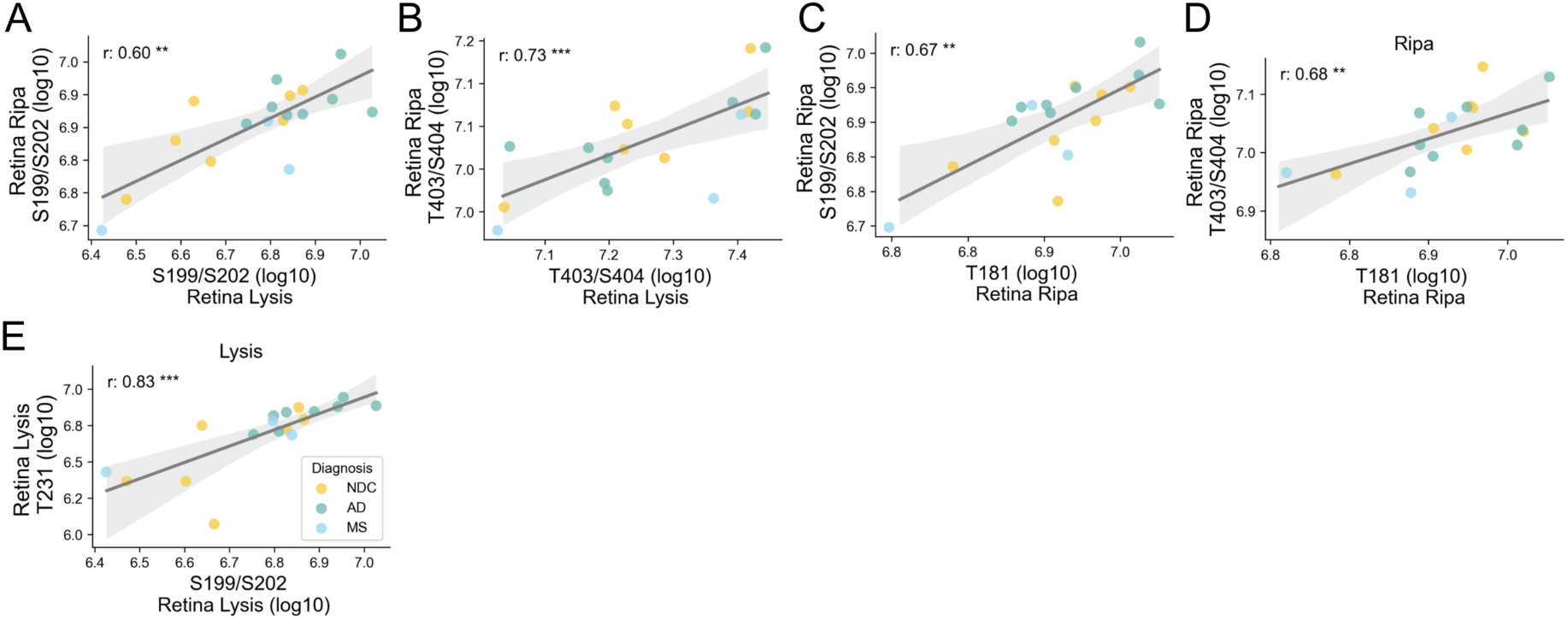
Correlation analysis of p-tau variants in the retina. Scatter plots showing significant correlations between retinal tau peptide phosphorylation peak intensities measured in Lysis and Ripa fractions. Across buffers, positive correlations were found between p-tau S199/S202 peptides in Ripa buffer and p-tau S199/S202 peptides in Lysis buffer (A) and between p-tau T403/S404 peptides in Ripa Buffer and p-tau T403/S404 peptides in Lysis buffer (B). Additionally, in the Ripa buffer, the peak intensities of p-tau T181 peptides correlated positively with p-tau S199/S202 peptides (C) and p-tau T403/S404 peptides (D). In the Lysis buffer, p-tau S199/S202 peptides showed a significant correlation with p-tau T231 peptides. Statistical analyses were performed using Spearman’s correlation tests. Each point represents an individual sample. Statistical significance is indicated by asterisks (*p < 0.05, **p < 0.01, ***p < 0.001).

The differences between AD and NC for the peptides S199/S202 (q=0,002), T231 (q=0.001) and total p-tau (q≤0.001) in the retina remained significant after multiple testing correction using Benjamin-Hochberg. Also, the differences between Aβ high and low groups for the retinal peptides phosphorylated at S199/S202 (q<0.001), T231(q=0.004), S396+T403/S404 total (q<0.001) and total p-tau (q<0.001) remained significant. In CA1, total p-tau (q=0.01) was still lower in MS compared to AD, and total p-tau was higher in AD than NC (q=0.01). The Aβ high group also showed higher values of S396+T403/S404 total (q<0.001) and total p-tau (q=0.04) in the brain. For the correlations with neuropathological stages of AD, only S396+T403/S404 with Aβ Braak Stages survived correction (q<0.001). Although none of the correlations between p-tau peptides in the retina and CA1 survived multiple testing correction, likely due to the large number of peptides tested in combination with the small sample size, the strong associations observed between S396+T403/S404 total, pathological staging, and multiple peptides in the hippocampus support the robustness of this finding. For the correlations within the retina, the associations of T181 with S199/S202 (q=0.001) and T403/S404 (q=0.002) in Ripa remained significant, as well as S199/S202 with T231 in Lysis (p<0.001) and the correlation of the values of T403/S404 between buffer fractions (p<0.001).

## DISCUSSION

While mass spectrometry has previously been successfully used to identify and quantify p-tau variants in the brain [7, 34] and cerebrospinal fluid [6, 40, 43], to our knowledge, no studies have yet applied this method to analyse p-tau in the human retina. The literature can therefore not directly support our identification and quantification of the six retinal tau peptides with phosphorylated sites at T181, S199/S202, T231, T231+T235, S396+T403/S404, and T403/S404. There are however several studies that have identified the same phosphorylation sites of tau in the retina using immunohistochemistry with antibodies directed against single p-tau sites (T181, S199, T231, and S396) [26, 49] or antibodies directed against double p-tau sites (AT8, directed against p-tau S202+T205 and PFH-1, directed against S396+S404) [16, 38, 49]. Notably, although previous immunostaining studies also have shown the presence of p-tau T212/S214 and T217 in the human retina [16, 26], we were unable to confirm this in the current study. This result may suggest that these p-tau variants are absent in the human retina; however, we believe it is more likely due to the abundance of the peptides which may be too low to be detected using our method. In support, previous work on retinal p-tau T217 has shown that the immunoreactivity of this p-tau variant is relatively much lower compared to other more physiologically relevant p-tau variants, such as p-tau T181[26]. This is however not the case with T212/S214 [26].

Our immunostainings of retina flat mounts showed a high abundance of scattered dots in the retina of NC when staining against p-tau at T181, T231, S202, S404, and 396+S404. The immunoreactivity strength varied, where staining against S404 yielded the strongest and 396+404 the lowest staining immunoreactivity. This result coincided by and large with the results found when we analysed the relative distribution (i.e. S404 yielded the highest, and 396+404 the lowest peak intensities; Figure 3). However, it is important to point out that the very high peak intensity value of the S404 peptide does not necessarily mean that this peptide is the most abundant, as values measured by mass spectrometry are subject to differences in ionization for different peptides. Nevertheless, based on these observations, we speculate that phosphorylation of S199/S202, T231, and S403/S404, just like T181, have a physiological role in the retina, while phosphorylation at S396+S403/S404 (detected foremost in cases with high Aβ Braak stages), in similarity to T217, arise foremost during pathological conditions. The notion that paired helical filaments (PHF) in the AD brain consist of tau phosphorylated at S396+S404 supports the idea that this p-tau variant is foremost pathological [39].

Our mass spectrometry study on retinal samples further revealed higher peak intensities of total p-tau peptides and peptides phosphorylated at S199/S202 and T231 in cases with AD pathology. No difference in p-tau T181 peptides was found, but higher peak intensities of p-tau S396+S403/S404 and total p-tau S403/S404 peptides were detected in retinas of individuals with higher stages of Aβ accumulation. These results are in line with previous immunohistochemistry and Nanostring GeoMx studies, which revealed higher p-tau T231, S404, S396+S404, and S202+T205 immunoreactivity in the retina of AD compared to control [49], while p-tau T181 appears to be less affected in the AD retina [26]. Of note, our immunostainings of the retina flatmounts did indicate a change in the immunoreactivity pattern (from dotted to striped staining pattern) of all p-tau variants in the AD case, including p-tau T181. This suggests that, although a potential increase in p-tau peak intensity is not detected, AD pathology may still contribute to intracellular alterations of p-tau.

Interestingly, while peptides with p-tau T231 and S199/S202 showed higher peak intensities in AD cases compared to NC in both CA1 and the retina, the pattern for MS cases differed between the tissues. In MS, the CA1 peak intensities resembled those seen in AD (indicating tauopathy also in the MS brain), whereas the retina intensities were closer to NC values (indicating no tauopathy in the MS retina). This finding suggests that retinal p-tau may be more specific to AD or classical tauopathies [26] and less affected in pathologies where p-tau accumulation may not be a primary event. Despite this, positive correlations were observed between some retinal p-tau peptides (S396+T403/S404, total T403/S404, and S199/S202) and NFT, LB, or Aβ Braak stages even when MS cases were included in the analysis. Similar correlations have been presented before using immunohistochemistry, where p-tau S202+S205, S396+S404, and S396 immunoreactivity in the retina were shown to correlate with NFT Braak stages [26, 49]. Another recent study also identified a correlation between p-tau S396 and both brain pathology and disease stage, as well as cognitive status and retinal ganglion cell loss[14]. Important to notice, S396+T403/S404 in the hippocampus of MS cases showed lower values than in AD, contrary to the other −tau peptides, indicating this phosphorylation site might be more AD specific in both tissue types. Our study further showed that the sum of retinal tau peptides phosphorylated at 396+T403/S404, correlated positively with their respective peptides in the CA1. Besides this correlation, we also noted that p-tau S396+T403/S404 correlated with several other p-tau variants in the CA1. Hence, although our study does not provide direct evidence that the retina is involved in a hierarchical spreading pattern of tau pathology, it does suggest that the phosphorylation patterns of tau, in particular S396+T403/S404, in the retina are linked to the phosphorylation of tau in the brain.

Lastly, one of the major advances of mass spectrometry compared to other assays such as immunohistochemistry, is its capability to compare peptides with single and double phosphorylation sites between different diagnostic groups. This capability can reveal specific temporal phosphorylation events linked to disease pathology. To exemplify, in NC cases the relative distribution of retinal p-tau variants for the most part followed the same pattern as the p-tau distribution in CA1, regardless of buffer condition. However, in AD cases, the peak intensity of peptides with a single p-tau at T231 was almost completely lost in the Ripa buffer (contains foremost organelles and nuclei), while peptides with the double p-tau at T231+T235 increased. The opposite pattern was seen in the Lysis buffer (containing foremost extracellular and cytosolic components), where instead MS showed the highest peak intensity of T231+T235. It is thus tempting to speculate that phosphorylation at T235 is an AD-related event occurring downstream of the T231 phosphorylation in the nuclei, while the phosphorylation at T235 in the cytosol/extracellular compartment is MS-related. The significance of such an event is yet to be determined, but the existence and a specific physiological function of p-tau in the nucleus have been suggested before [17, 20, 53]. Indeed, in our study, only p-tau S199/S202 and T403/S404 in the Lysis buffer correlated with the same in the Ripa buffer, which again suggests that some p-tau variants might behave differently in the different cellular compartments.

### Limitations

This study provides valuable insights into retinal tau pathology; however, some limitations must be acknowledged. The sample size was limited for each diagnostic group, reducing statistical power and the generalizability of findings. Moreover, analysis of postmortem tissue do not fully represent in vivo conditions and the dynamic changes in tau phosphorylation. Detection sensitivity posed another challenge, as certain phosphorylation sites previously found in the retina were not identified, and others were identified in less than 50% of subjects, likely due to low levels of the peptide in the sample, though technical limitations may also have contributed. Furthermore, due to limited access to tissue, we were only able to analyse a part of the middle periphery of the superior nasal retina. Examining other retinal regions could potentially reveal other phosphorylation patterns, as previous studies have shown that p-tau is predominantly localized in the far periphery of the retina[16, 26].

## Conclusion

By employing mass spectrometry analysis of retina and CA1 samples from NC, AD, and MS cases, we have identified the most abundant p-tau variants in the retina. This analysis has also demonstrated that some of the abundant p-tau variants, more specifically, T231, S199/S202 and 396+T403/S404 are significantly more common in cases with Aβ pathology. Among these, p-tau 396+T403/S404 stands out as the variant most strongly associated with Braak Aβ and NFT stages, as well as with p-tau variants in CA1. These findings support the hypothesis that the retina may serve as a valuable source of biomarkers for neurodegenerative diseases, particularly for AD diagnosis and progression monitoring, while also advancing our understanding of tauopathies in both the retina and the brain.

## DECLARATIONS

### Ethics approval and consent to participate

Informed consent for using retina and brain tissue as well as clinical data for research purposes was obtained from the patients or from their closest relatives in accordance with the International Declaration of Helsinki and the Code of Conduct for Brain Banking. The tissue collection protocols were approved by the medical ethics committee of VU Amsterdam and the Swedish Ethical Review Authority approved the study (Dnr 2021/04270). All data were analysed anonymously.

### Consent for publication

Not applicable.

### Availability of data and material

The data supporting the findings of this study are available from the corresponding author upon reasonable request.

### Authors’ contributions

J.S and M.W contributed to the study concept and design. D.P prepaired the samples. J.S analysed the data with support from J.V and was a major contributor in writing the manuscript. The NBB contributed to the brain tissue and neuropathological evaluation. G.B determined the phosphorylation sites on the peptides with PEAKS Studio. All authors have read and revised the manuscript for intellectual content and agreed to the final version of the manuscript.

## Acknowledgments

The authors wish to thank the Center for Translational Proteomics at the Medical Faculty, Lund University for the proteomic analysis. The authors also wish to thank Dr. Charlotte Wehlinder, Dr. Patrik Önnerfjord, and Dr. Nina Schultz for valuable discussion and scientific input.

## Abbreviations

2N4R: Tau isoform with two amino-terminal inserts and four microtubule-binding repeats
ACN: Acetonitrile
AGC: Automatic Gain Control
Aβ: Amyloid Beta
AD: Alzheimer’s Disease
CA1: Cornu Ammonis 1 (hippocampal region)
DAPI: 4′,6-diamidino-2-phenylindole
DDA: Data-Dependent Acquisition
DNA: Deoxyribonucleic Acid
DTT: Dithiothreitol
EDTA: Ethylenediaminetetraacetic Acid
FA: Formic Acid
HCD: Higher-Energy Collisional Dissociation
HT7: Total Tau Antibody Clone
IgG: Immunoglobulin G
LB: Lewy bodies
LC-MS/MS: Liquid Chromatography–Tandem Mass Spectrometry
m/z: Mass-to-Charge Ratio
MS: Multiple Sclerosis
NC: Normal Control
NBB: The Netherlands Brain Bank
NFTs: Neurofibrillary Tangles
PHFs: Paired Helical Filaments
PBS: Phosphate-buffered saline
PMD: Post-Mortem Delay
PTMs: Post-Translational Modifications
p-tau: Phosphorylated Tau
RD3: 3-Repeat Tau Antibody Clone
RD4: 4-Repeat Tau Antibody Clone
RT: Room Temperature
S404: Serine 404
T181: Threonine 181
T231: Threonine 231
TAR: Transactive Response DNA-binding protein
TFA: Trifluoroacetic Acid
UHPLC: Ultra High-Performance Liquid Chromatography
MAPT: Microtubule-Associated Protein Tau gene
mRNA: Messenger Ribonucleic Acid

