## Supplement for "Retinal tau phosphorylation in Alzheimer’s disease: a mass spectrometry study"

**Supplement Table S1**. Demographic data and neuropathological evaluation of the cases included in the study.

| Diagnosis | Age  (years) | Biological sex | NFT (Braak) | Aβ  (Braak) | LB  (Braak) | *APOE*  alleles | PMD  (h:m) |
| --- | --- | --- | --- | --- | --- | --- | --- |
| NDC^r,b^ | 75 | male | I | A | 0 | 33 | 05:25 |
| NDC^r,b^ | 81 | male | III | C | 0 | 43 | 04:30 |
| NDC^r,b^ | 92 | female | III | O | 1 | 34 | 06:35 |
| NDC^r,b^ | 68 | female | I | O | 0 | 33 | 04:30 |
| NDC^r,b^ | 84 | female | II | B | 0 | 33 | 06:05 |
| NDC^r,b^ | 102 | male | III | A | 0 | 43 | 05:00 |
| NDC^b^ | 60 | female | 0 | O | 0 | 32 | 08:10 |
| NDC^r,b^ | 70 | male | I | O | 3 | 32 | 06:20 |
| AD^r,b^ | 92 | female | VI | C | 0 | 43 | 06:10 |
| AD^b^ | 78 | female | VI | C | 0 | 44 | 04:45 |
| AD^r,b^ | 70 | female | VI | C | 0 | 44 | 04:20 |
| AD^b^ | 85 | male | IV | C | 0 | 33 | 08:35 |
| AD^r,b^ | 65 | female | V | C | 0 | 43 | 04:30 |
| AD^r,b^ | 88 | female | V | C | 0 | 43 | 08:15 |
| AD^r,b^ | 83 | female | IV | C | 6 | 33 | 06:15 |
| AD^r,b^ | 63 | male | IV | C | 6 | 44 | 04:55 |
| AD^b^ | 88 | female | V | C | 5 | 43 | 05:40 |
| AD^b^ | 91 | female | IV | C | 0 | 33 | 06:40 |
| AD^r,b^ | 96 | female | IV | B | 0 | 33 | 07:20 |
| AD^r,b^ | 69 | male | VI | C | 0 | 43 | 06:30 |
| MS^r,b^ | 60 | female | 0 | O | 0 | 43 | 07:25 |
| MS^r,b^ | 78 | male | 0 | O | 0 | 33 | 08:45 |
| MS^r,b^ | 81 | female | IV | O | 0 | 33 | 09:35 |
| MS^b^ | 87 | female | II | O | 0 | 33 | 09:30 |

NDC=non-demented controls, AD=Alzheimer’s disease, NFT = neurofibrillary tangles, Aβ = amyloid beta, LB = Lewy bodies, *APOE* = apolipoprotein E, PMD= post-mortem delay. ^r^ Included in the analysis of the retina. ^b^ Included in the analysis of the brain.

**Supplement Table S2.** Differences in p-tau levels in the retina across diagnostic groups and Aβ high vs low groups.

| p-tau site | Comparison | 95% CI | p-value | q-value |
| --- | --- | --- | --- | --- |
| S199/S202 | AD vs NC | 0.064 – 0.281 | 0.0019 | 0.0186 |
| T231 | AD vs NC | 0.072 – 0.463 | 0.0074 | 0.0491 |
| T231 | AD × Ripa interaction | -0.799 – -0.406 | 1.9e-09 | 7.6e-08 |
| T231 total | MS × Ripa interaction | -0.911 – -0.142 | 0.0072 | 0.0491 |
| Total phosphorylation | AD vs NC | 0.051 – 0.175 | 0.00037 | 0.0073 |
| Total phosphorylation | AD × Ripa interaction | -0.201 – -0.054 | 0.00071 | 0.0094 |
| S199/S202 | Aβ high vs low | 0.068 – 0.264 | 0.00088 | 0.0035 |
| T231 | Aβ high vs low | 0.088 – 0.472 | 0.0042 | 0.0141 |
| T231 | Aβ high × Ripa interaction | -0.675 – -0.186 | 0.00056 | 0.0035 |
| T231+T235 | Aβ high vs low | -1.185 – -0.197 | 0.0061 | 0.0174 |
| S396+T403/S404 | Aβ high vs low | 0.224 – 0.720 | 0.00019 | 0.0035 |
| S396+T403/S404 total | Aβ high vs low | 0.206 – 0.721 | 0.00042 | 0.0035 |
| Total phosphorylation | Aβ high vs low | 0.040 – 0.154 | 0.00080 | 0.0035 |

| After adjustment for age, sex, and postmortem delay: | | | | |
| --- | --- | --- | --- | --- |
| S199/S202 | AD vs NC | 0.063 – 0.281 | 0.0045 | 0.0455 |
| T231 | AD × Ripa interaction | -0.798 – -0.406 | 1.9e-09 | 7.6e-08 |
| Total phosphorylation | AD vs NC | 0.051 – 0.175 | 0.0031 | 0.0408 |
| Total phosphorylation | AD × Ripa interaction | -0.201 – -0.054 | 0.00071 | 0.0251 |
| S199/S202 | Aβ high vs low | 0.068 – 0.264 | 0.0128 | 0.0492 |
| T231 | Aβ high vs low | 0.088 – 0.472 | 0.0148 | 0.0492 |
| T231 | Aβ high × Ripa interaction | -0.676 – -0.186 | 0.00056 | 0.0056 |
| S396+T403/S404 | Aβ high vs low | 0.505 – 0.721 | 0.00030 | 0.0056 |
| S396+T403/S404 total | Aβ high vs low | 0.471 – 0.721 | 0.0017 | 0.0087 |
| Total phosphorylation | Aβ high vs low | 0.041 – 0.154 | 0.0010 | 0.0068 |

Only results with false discovery rate–corrected p-values (q-values) < 0.05 are included. Two buffer types were used for protein extraction: Lysis (reference) and Ripa. Group comparisons are fixed with non-demented controls (NC) or Aβ-low individuals as the reference group. Results are shown both before and after adjustment for age, sex, and postmortem delay. Values represent pairwise group differences, with 95% confidence intervals (CI), p-values, and FDR-corrected p-values (q-values).

**Supplement Table S3.** Differences in p-tau between diagnostic groups in the CA1 area of the hippocampus.

| p-tau site | Comparison | 95% CI | p-value | q-value |
| --- | --- | --- | --- | --- |
| S199/S202 | AD vs NC | 0.020 – 0.159 | 0.0117 | 0.039 |
| S199/S202 | MS vs NC | 0.099 – 0.286 | 0.000056 | 0.0011 |
| S199/S202 | MS × Ripa interaction | -0.186 – -0.037 | 0.0034 | 0.0167 |
| T231 | AD vs NC | 0.021 – 0.087 | 0.0013 | 0.0088 |
| T231 | MS vs NC | 0.011 – 0.100 | 0.0140 | 0.0430 |
| T231+T235 | AD vs NC | 0.055 – 0.251 | 0.0022 | 0.0124 |
| S396+T403/S404 | AD vs NC | 0.024 – 0.228 | 0.0157 | 0.0449 |
| T403/S404 | MS vs NC | 0.050 – 0.358 | 0.0096 | 0.0349 |
| T231 total | AD vs NC | 0.041 – 0.213 | 0.0038 | 0.0167 |
| T403/S404 total | AD vs NC | 0.056 – 0.223 | 0.0010 | 0.0088 |
| T403/S404 total | AD × Ripa interaction | 0.038 – 0.157 | 0.0013 | 0.0088 |
| S396+T403/S404 total | AD vs NC | 0.115 – 0.433 | 0.0007 | 0.0088 |
| Total phosphorilation | AD vs NC | 0.052 – 0.149 | 0.000051 | 0.0011 |
| Total phosphorilation | AD × Ripa interaction | 0.015 – 0.086 | 0.0050 | 0.0201 |

| After adjustment for age, sex, and post mortem delay: | | | | |
| --- | --- | --- | --- | --- |
| S199/S202 | MS × Ripa interaction | –0.186 – –0.037 | 0.0034 | 0.0201 |
| T231 | AD vs NC | 0.023 – 0.091 | 0.0009 | 0.0174 |
| T231+T235 | AD vs NC | 0.047 – 0.238 | 0.0035 | 0.0201 |
| T231 total | AD vs NC | 0.033 – 0.204 | 0.0064 | 0.0283 |
| T403/S404 total | AD vs NC | 0.049 – 0.233 | 0.0027 | 0.0201 |
| T403/S404 total | AD × Ripa interaction | 0.038 – 0.157 | 0.0013 | 0.0176 |
| S396+T403/S404 total | AD vs NC | 0.100 – 0.437 | 0.0018 | 0.0179 |
| Total phosphorilation | AD vs NC | 0.047 – 0.154 | 0.0003 | 0.0102 |
| Total phosphorilation | AD × Ripa interaction | 0.015 – 0.086 | 0.0050 | 0.0251 |

Only phosphorylated peptides that are also present in the retina and results with false discovery rate–corrected p-values (q-values) < 0.05 are included. Two buffer types were used for protein extraction: Lysis (reference) and Ripa. Group comparisons are fixed with non demented controls (NC) as the reference group. Results are shown both before and after adjustment for age, sex, and postmortem delay. Values represent pairwise group differences, with 95% confidence intervals (CI), p-values, and FDR-corrected p-values (q-values).

**Supplement Table S4.** Associations between retinal phospho-tau levels and Aβ, neurofibrillary tangle (NFT), and Lewy body (LB) pathology stages.

| p-tau site | Beta | p-value | q-value |
| --- | --- | --- | --- |
| **Associations with Aβ stages** | | | |
| S396+T403/S404 total (lysis) | 0.182969 | 0.003142 | 0.025135 |
| Total phospho (lysis) | 0.040624 | 0.031819 | 0.101460 |
| T403/S404 (lysis) | 0.050954 | 0.047057 | 0.101460 |
| T231 (lysis) | 0.085264 | 0.050730 | 0.101460 |
| **Associations with NFT stages** | | | |
| S396+T403/S404 total (lysis) | 0.099957 | 0.025342 | 0.171212 |
| Total phospho (lysis) | 0.025230 | 0.051521 | 0.171212 |
| T403/S404 (lysis) | 0.032336 | 0.064204 | 0.171212 |
| **Associations with LB stages** | | | |
| S396+T403/S404 total (lysis) | 0.084283 | 0.024363 | 0.194903 |

Linear regression models were used to assess the associations between retinal phospho-tau markers and Aβ, NFT, and LB pathology stages, adjusting for age, sex, and postmortem delay. Only associations with raw p-values < 0.065 are shown. FDR correction was applied using the Benjamini-Hochberg method.

**Supplement Table S5**. Associations between retinal phospho-tau levels at different sites and buffers.

| **Predictor** | **Outcome** | **Beta** | **p-value** | **q-value** |
| --- | --- | --- | --- | --- |
| **Ripa x Ripa** | | | | |
| T403/S404 total | S396+T403/S404 total | 2.1021 | 0.0000 | 0.0000 |
| T181 | S199/S202 | 0.8656 | 0.0086 | 0.0775 |
| T181 | T403/S404 | 0.9211 | 0.0120 | 0.0845 |

| **Lysis x Lysis** | | | | |
| --- | --- | --- | --- | --- |
| S199/S202 | T231 | 1.0923 | 0.0004 | 0.0151 |

| **Lysis x Ripa** | | | | |
| --- | --- | --- | --- | --- |
| T403/S404 | T403/S404 | 0.4028 | 0.0000 | 0.0014 |
| T231 | S199/S202 | 0.2254 | 0.0000 | 0.0014 |
| S199/S202 | S199/S202 | 0.2788 | 0.0006 | 0.0151 |
| T231 | T181 | 0.1372 | 0.0034 | 0.0543 |
| S199/S202 | T181 | 0.1664 | 0.0137 | 0.0845 |

Linear regression models were used to assess the associations between different retinal p-tau sites and buffers, adjusting for age, sex, and postmortem delay. Only associations with raw p-values < 0.05 are shown. FDR correction was applied using the Benjamini-Hochberg method
